# Model-based separation of visual and oculomotor signals in marmoset lateral intraparietal area

**DOI:** 10.64898/2025.12.01.691532

**Authors:** Daisuke Shimaoka, Joanita F. D’Souza, Maureen A. Hagan, Nicholas S. C. Price

## Abstract

The lateral intraparietal area (LIP) integrates visual information with task demands to guide saccadic eye movements. Most prior studies have relied on imposed prolonged delays to separate sensory and motor activity, whereas natural behavior involves rapid sensory-motor transformations. To examine LIP function during more naturalistic oculomotor behavior, we recorded multi-unit activity from LIP in two marmosets performing a task requiring rapid selection of salient visual stimuli. Because sensory and motor signals temporally overlap in such fast tasks, we used a Generalized Linear Model (GLM) that separately captured visually driven and oculomotor-related components. The model explained substantial neural variance during correct and missed saccades, as well as spontaneous eye movements, and revealed heterogeneous sensory and motor encoding across recorded neural units. Our approach demonstrated that the origins of neural activity can be segregated without relying on imposed delays, and showed the presence of a strong post-saccadic motor signal in LIP.

## Introduction

The posterior parietal cortex (PPC) plays a crucial role in linking sensory input to motor output, supporting functions such as spatial attention, decision-making, and motor planning^1^. Within PPC, the lateral intraparietal area (LIP) is particularly implicated in integrating visual signals with task demands to guide saccadic eye movements^2,3^. Anatomically, LIP is well positioned within the frontoparietal network, possessing reciprocal monosynaptic connections with areas involved in visual and motor processing, including the frontal eye fields (FEF) and superior colliculus (SC)^4,5^. Despite this central role, the functional diversity across cells within LIP and how distinct neuronal populations contribute to sensory-motor transformations are not yet fully understood.

Many neurons within LIP exhibit robust pre-saccadic firing^6,7^ and electrical stimulation of LIP reliably evokes saccades of a particular direction and amplitude in macaques^8,9^ and marmosets^10^. However, saccades induced by electrical stimulation have relatively long and variable latencies (> 50 ms) with stronger currents (up to 250 µA in marmosets^10^), compared to latencies induced at FEF^11^. This suggests that LIP is not normally involved in the generation of commands of saccade execution, but can influence saccade probability and modulate its parameters^12,13^. Reversible inactivation of LIP leads to longer saccade latencies, particularly for tasks using memory-guided saccades, multiple targets, and covert visual search tasks^14,15^. Beyond motor planning, neuronal activity in LIP is influenced by cognitive variables such as attention^16,17^, reward^18^, decision-making^19–21^ and working memory^6,22^, supporting theories that LIP acts as a dynamic “priority map” integrating top-down executive signals with bottom-up sensory inputs, to guide behavior^23,24^.

Electrophysiological studies have revealed functionally heterogeneous LIP subpopulations, responsive to sensory processing (visual stimulus onset), memory-related activity during delay periods, and visuomotor neurons encoding saccade planning and execution^6,7,25–27^. The functional diversity across LIP suggests that neurons here support multiple stages of sensory-motor processing, from stimulus detection to motor execution. However, most previous work has relied on prolonged delay periods to temporally dissociate sensory and motor signals^7,28^. This contrasts natural behavior, in which short-latency saccades to salient targets arise from rapid sensory-motor transformations^29^. As a result, there remains a substantial gap in our understanding of how distinct LIP neuron classes contribute to rapid, real-time sensory-motor computations.

To investigate how LIP contributes to natural, routinely performed oculomotor behavior, we used a task that requires rapid selection of salient and behaviorally relevant visual stimuli. In such fast, goal-directed tasks, visual sensory representations are expected to be expressed concurrently with oculomotor signals. Unlike behavioral tasks that impose long delay periods, this design makes it more challenging to dissociate whether neural activity reflects sensory processing, motor planning, or a mixture of both.

To address this challenge, we adopted a computational modeling approach to quantitatively characterize and predict neural firing. Specifically, we employed a Generalized Linear Model (GLM)^30^ that includes distinct components for visually-driven responses and oculomotor-related activity to multi-unit electrophysiological activity recorded from neural units in LIP. By fitting this model to spiking data from individual neural units in LIP, we were able to decompose the observed firing patterns into their contributing components. Our modelling builds on previous successful applications of GLMs to visual discrimination and memory-guided oculomotor tasks in LIP and FEF^31–34^, extending it to the relatively understudied domain of rapid, goal-directed behavior.

Critically, if saccades always occurred with fixed latency after a visual stimulus, it would be impossible to distinguish sensory from motor activity. Natural variations in reaction time helped address this, and we also considered responses to spontaneous saccades that were not driven by our visual stimuli, and visual stimuli that were not followed by saccades. Through this decomposition, we aimed to address two key questions: 1) what role do individual LIP neurons play in the oculomotor transformation; and 2) how do individual LIP neurons contribute to the initiation of saccades?

Our analyses based on GLM successfully identified visually responsive and oculomotor-responsive neural units, consistent with prior reports^35^. Neural units that appeared to dissociate correct from failed stimulus detection could almost entirely be explained by their oculomotor-related activity. Moreover, motor-related units typically began responding only after saccade onset. Together, these findings suggest that while marmoset LIP robustly participates in rapid, naturalistic oculomotor behavior, its role may lie downstream of detection of visual stimuli and motor command initiation, highlighting a more nuanced function in the sensory-motor transformation process.

## Methods

Two adult male marmoset monkeys (*Callithrix Jacchus*) served as subjects in this study (M1, 6 years old, 440g; M2, 4 years old, 490g). All procedures were approved by the Monash Animal Research Platform Animal Ethics Committee and followed the Australian Code of practice for the care and use of animals for scientific purposes.

### Surgical procedure

Prior to training on the behavioral task, marmosets were implanted with a titanium head-post to stabilize the head during experiments, and a titanium cranial chamber over LIP (NeuroNexus, USA)^36^. The marmosets were first injected with atropine (0.2 mg/kg, i.m.) and diazepam (2 mg/kg, i.m.). After 30 minutes, anesthesia was induced with alfaxalone (8 mg/kg, i.m. Jurox, Rutherford, Australia). The marmoset was then placed in a stereotaxic frame and stabilized using earbars, which had been covered in local anesthetic (2% xylocaine jelly). After intubation, the head was further stabilized with a palate bar and eye bars. Anesthesia was maintained by isoflurane (0.5-3%) in oxygen. Eyes were protected during surgery with liquid paraffin eye ointment. Once stabilized, a midline incision was made, the scalp was reflected and the temporalis muscles separated to expose the skull. Up to six titanium screws (diameter 1.5 mm, length 4 mm) were inserted 1-1.5 mm into the skull. The exposed surface of the skull was coated with a thin layer of dental adhesive (Suparbond, Sunmedical, Japan). A head-post, which would stabilize the animal’s head during experiments, was then placed on one hemisphere, as close to the midline as possible, with a transparent dental acrylic (Ortho-Jet; Lang Dental Mfg. Co., USA). The margin was cleaned and the skin sealed with surgical adhesive (VetBond; 3M, USA) to the base of the headcap. Animals received oral antibiotics for 7 days (cefalexin monohydrate, 30 mg/kg) and analgesia for 5 days (meloxicam, 0.2 mg/kg). In a separate surgery for M1, but the same surgery for M2, a 9 mm diameter craniotomy, which left the dura intact, was performed in one hemisphere over LIP at stereotaxic coordinates anterior-posterior 0, medial-lateral 6 mm^37,38^. The cranial chamber was then placed over this region and secured in place on the skull with the dental and dental acrylic. A postmortem histological examination of M1 indicated the region is densely myelinated, an indicator of marmoset LIP^39^.

### Behavioral training

One week after surgery, the marmosets were acclimated to sitting in a custom designed primate chair with their head position fixed. Visual stimuli were presented on a 24-inch Viewpixx/3D LCD monitor (VPixx Technologies Inc., Canada) with a resolution of 1920 × 1080 pixels (W × H), and a refresh rate of 100 Hz. The stimulus monitor was positioned at a fixed viewing distance of 48 cm in front of the marmoset. All visual stimuli were generated in MATLAB (MathWorks, Inc, USA) using Neurostim (https://klabhub.github.io/neurostim/) and the Psychophysics Toolbox extension^40^. Eye position was tracked monocularly with a video-based Eyelink 1000 system (SR Research, UK). Horizontal and vertical eye positions of the right eye were recorded at 1 kHz sampling rate. At the start of each session, eye positions were calibrated by presenting marmoset faces (3.5 degrees visual angle) at different locations in the visual field, up to 10 degrees eccentricity. The marmosets were rewarded with sweetened liquids such as fruit juice (up to 12 mL in a session) at the end of every correctly completed trial (approximately 0.035 mL a trial, through a New Era syringe pump system).

### Center-out saccade task design

The center-out saccade task (**Figure 1A**), required marmosets to first centrally fixate within a 2 degrees radius window (700 - 900 ms for M1; 500 - 700 ms for M2) to initiate a trial.

**Figure 1.**
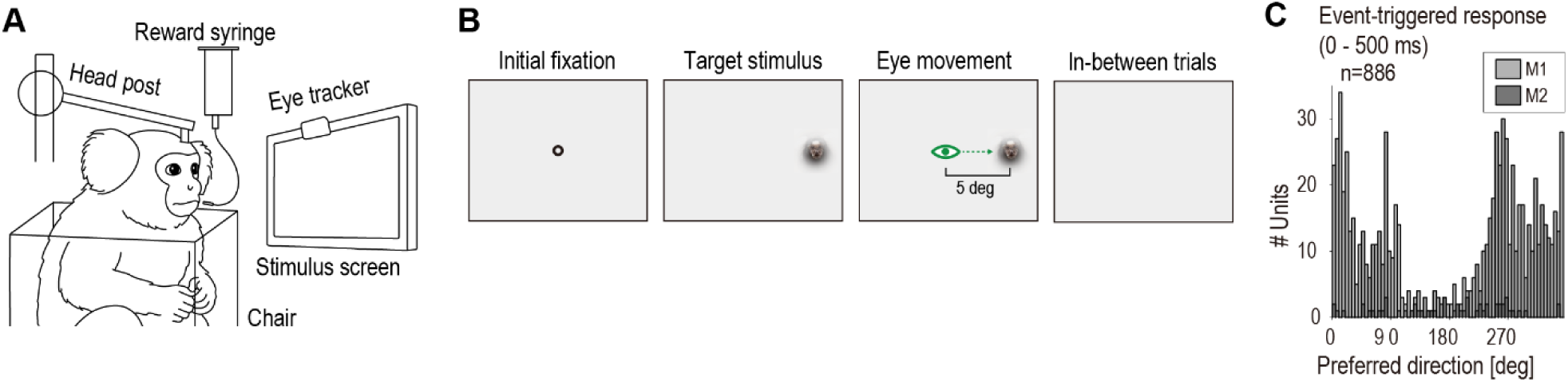
Center out-saccade task. **A**. Animal recording setup. **B**. Task trials started with presentation of a fixation point, followed by presentation of visual stimulation. The animal was trained to make a saccadic eye movement to the target direction. In between trials, the animal kept spontaneous, untrained eye movements. **C**. Distribution of preferred direction across recorded units, from two animals, M1 and M2.

Thereafter, a marmoset face (2.5° visual angle) appeared at 5° eccentricity. On different days, these peripheral saccade targets could appear at one of either 8 or 36 equidistant locations. Marmosets had to saccade to the peripheral target stimulus within 500 ms and hold fixation within a two degrees radius tolerance window for 200 - 400 ms to correctly complete a trial and receive a juice reward. Mean (SD) reaction times for the two animals were: M1: 124.2 (59.0) ms; M2: 159.5 (91.2) ms. Marmosets completed an average of 280 trials/session, with sessions lasting one to two hours.

While analyses in this paper were only conducted on trials without any spatial cueing, a subset of trials (35%, randomly interleaved) included a spatial cue. This cue consisted of a small grey bar that appeared 200 ms after fixation onset for 50 ms, at 1.5° eccentricity, and indicated the location of the upcoming target with 85% accuracy. The cue’s location remained fixed at a specific direction within a single recording session. Positioned at a fixed location outside the fixation window, the cue required the marmosets to attend covertly, and trials in which they made direct saccades to it were immediately aborted. Neither animal showed a significant difference in saccade reaction times between cued and uncued trials at the cued location (M1: U = 20869,062, p = 0.05, cued median = 115 ms, uncued median 116 ms; M2: U = 591.537, p = 0.08, cued median = 136.5 ms, uncued median = 142.2 ms, Mann-Whitney U test)

### Electrophysiological recordings

Once a marmoset was reliably performing the center-out saccade task to at least a 70% completion rate, a semi-chronic NeuroNexus p-drive housing a 32-electrode laminar probe (100 µm electrode spacing) was implanted through the recording chamber for two to six weeks. The structure of the micro-drive allowed us to position the electrode throughout the recording chamber, constrained by a radial grid^36^. The external reference and ground wires were bridged and grounded to a silver wire that rested on the dural surface. Once the array had penetrated the dura and ground wire was in position, the recording chamber was filled with duragel (Cambridge NeuroTech, UK).

In total, we collected data in 279 recording sessions from 10 array penetrations in M1 (left hemisphere) and 8 penetrations in M2 (right hemisphere). In total, we isolated 3784 units in these sessions. Electrodes were mounted to a drive screw that allowed us to advance the electrode over the course of the implant as needed to isolate cells. Two drive designs were used, with each turn of the drive screw advancing the array 150 or 250 µm, with a total drive range of 7 or 12 mm. Typically, on the day of insertion, we advanced the probe until we saw neural activity to depths of at least 400 µm. Thereafter, we would slowly advance 0 - 250 µm per day over the next 4 weeks until the array was fully implanted. As such, we were able to sample from all cortical depths in LIP.

Neural activity from each electrode was sampled at 30 kHz (OpenEphys, USA). Multiunit spiking activity was extracted from each channel by high-pass filtering the raw voltage traces at 300 Hz and applying a 5 standard deviation threshold to identify putative spikes.

### Unit inclusion criteria

In a fraction of recorded units, we observed an infrequent but exceedingly high spike rate. We removed trials that included periods of at least 20 ms in which the spike rate exceeded the median + 10*mad. On average across units from the two subjects, 3.3% of trials were excluded by this procedure. We subsequently selected units that met the following four criteria: 1) at least 200 trials in which the target stimulus was presented; 2) mean firing rate throughout the recording period was at least 5 spikes/s; 3) the unit shows modest direction sensitivity 0 - 500 ms after the onset of target stimuli, evaluated as p-value less than 0.2 with dot-product test; and 4) the encoding model provided a reasonably good fit, evaluated as correlation between observed and modelled response, averaged across correct trials, to be at least 0.5. Each of the four criteria was applied separately and only those that collectively meet each criterion were subject to further analysis.

### Encoding Model

We first created a spike density function (SDF) by counting spikes in 20 ms bins and then smoothing these counts with a half-Gaussian causal filter with 25 ms standard deviation. Finally, responses were detrended with high pass filtering with a cutoff of 0.001 Hz.

We employed a GLM to predict neural firing using five explanatory variables related to the visual stimulus and eye movements. Each explanatory variable is described in detail below.

#### 1) Target visual stimulus position

A matrix of 26 time points × 8 angular positions was defined, corresponding to the time period 0 - 500 ms after target stimulus onset, with 20 ms resolution, and 8 equally sized bins centered on 0, 45, 90, 135, 180, 225, 270, and 315° in which a stimulus could appear. Zero degree as defined as the horizontal direction to the subject’s right. In some sessions, there were 8 stimulus positions, which perfectly align with these bins, whereas in others there were 36 stimulus positions, so the bins contain 4 or 5 possible positions. Each bin was assigned a value of 1 or 0, representing the absence or presence of the target stimulus at that time and angle. This allowed us to model the visual receptive fields of neurons, which have been reported to tile the visual field^36^. The collective influence of visual stimulus position in the model is given by:

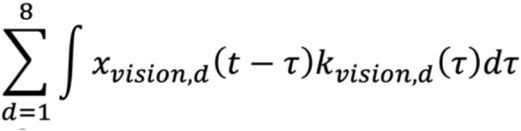

where ***x_vision_, d(t)*** represents the visual stimulus presented at a quantized direction ***d*** at time ***t***, ***k_vision_, d(t)*** represents the temporal kernel for the stimulus direction ***d*** at time ***t*** before stimulus presentation.

#### 2) Eye position relative to the fixation point

Similar to target position, we defined a matrix of 38 time points × 8 positions, corresponding to the time period from -250 to 500 ms relative to the time when eye position is recorded. Missing eye position data during blinks were linearly interpolated. Similar to target position, eye angular position relative to the fixation point was subdivided into eight angular bins. Each bin was assigned a position eccentricity value. These angular bins allowed representation of direction selectivity of eye-position signals, as reported previously^41,42^. For each angular bin, the period for -250 to 500 ms relative to the eye position change was modelled. This temporal window allowed modeling of eye-position related firing that has been observed before and after eye movements^43^. The collective influence of eye position in the model is given by:

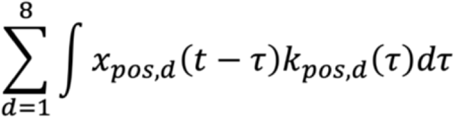

where ***x_pos, d_(t)*** represents the visual stimulus presented at a quantized direction ***d*** at time ***t***, ***k_pos, d_(t)*** represents the temporal kernel for the eye position direction ***d*** at time ***t*** before eye position event.

#### 3) Eye speed

Similar to eye position, we defined a matrix of 38 time points × 8 positions, corresponding to the time period from -250 to 500 ms relative to change in eye speed. Eye position data was smoothed using Savitzky-Golay Filtering (polynomial order 3 and frame length 11), then missing data during blinks were interpolated linearly. Velocity was computed as the temporal derivative of the filtered position, then converted into angular speed and eccentricity. For any given moment, the eccentricity of the eye speed was assigned to one of the eight bins that matches the quantized angular speed. This temporal window allowed modeling of both pre-saccadic firing^44–47^ as well as post-saccadic firing^35,48^. The number of modelled variables were 304, corresponding to 38 time points × 8 directions. The collective influence of eye speed in the model is given by:

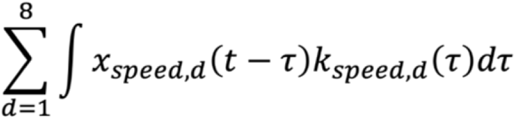

where ***x_speed, d_(t)*** represents the visual stimulus presented at a quantized direction ***d*** at time ***t***, ***k_speed, d_(t)*** represents the temporal kernel for the eye position direction ***d*** at time ***t*** before eye speed event.*Pupil diameter*. Changes in pupil diameter were modeled for -500 to 500 ms, corresponding to 51 modelled variables. This component is given by:

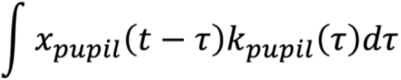

where ***x_pupil_(t)*** represents the pupil diameter at time ***t***, ***k_pupil_(t)*** represents the temporal kernel at time ***t*** before a pupil event.

#### 4) Eye blinks

Similar to pupil diameter, we defined a matrix of 51 time points, corresponding to the time period from -500 to 500 ms relative to the eye blink onset. Blinks were identified with an algorithm implemented in Eyelink. Periods 50 ms before and after the detected blinks were classified as periods of blinks in this study. Each moment in time was labelled with one (detected blinking) or zero (no detected blinking). This component is given by:

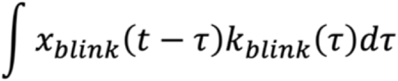

where ***x_blink_(t)*** represents the blink event at time ***t***, ***k_blink_(t)*** represents the temporal kernel at time ***t*** before a blink event.

In total, 918 variables were estimated in the GLM, corresponding to: (1) 26 time points × 8 directions (target position); (2) 38 time points × 8 directions (eye position); (3) 38 time points × 8 directions (eye velocity); (4) 51 time points (pupil diameter); and (5) 51 time points (blinks).

Five-fold cross-validation with ridge regression was used to fit the GLM. To assess the performance of the estimated model, Pearson correlation between the modeled and observed SDFs was computed. Modeling and evaluation procedures were repeated across five folds, with different partitions of the data used for modeling and evaluation, and the average GLM weights were used for further analysis.

To evaluate the performance of the individual linear kernels, we also computed the correlation between the observed SDF against a modeled SDF using only a subset of kernels. For example, to evaluate a vision only model, only the Target visual stimulus position kernel was used, with all other kernels set to 0. By taking the ratio of the correlation computed from the subset of kernels, using one out of the five kinds of explanatory variables, against the correlation from the full model using the five kinds of explanatory variables, we estimate the dominance of the subset of kernels in explaining observed SDFs. We compared the dominance amongst the three key explanatory variables, visual stimulus, eye speed and eye position.

### Direction Preference

The preferred direction of each unit was defined from the activity aligned to the onset of the target stimuli. For each stimulus direction, the mean firing rate in the 0 - 500 ms post-onset window was calculated, and baseline activity, averaged over -100 - 0 ms, was subtracted to obtain a direction-specific response amplitude. This triggered-average was then fitted with a Gaussian function wrapped at 0 and 360 degrees. Finally, the estimated center of the Gaussian was quantized and registered as the preferred direction.

### Neural Correlate of Task performance

Trials were classified as “correct” or “failed”. Correct trials were those in which animals correctly made an eye movement to the target position, followed by liquid reward. Failed trials were defined as the remainder of trials, in which the target stimulus was presented but the animals did not move their eyes to the target stimulus.

To compute how well the neuronal firing discriminated between correct and failed trials, we focused on trials where the target stimuli were presented at the unit’s preferred direction. Neuronal activity during these trials was summarized as the SDF averaged over the 30 - 250 ms window after target stimulus onset. In each unit, the discriminability of neuronal activity in correct versus failed trials was quantified through analysis of the Receiver-Operation Characteristic (ROC) curve. The area under ROC (AUC) for each unit indicated the strength of the correlation between the unit’s firing and the apparent stimulus-detection behavior.

Across the units, we examined distribution of the neural correlates of task performance, with respect to their encoding of different aspects of the task: vision, eye speed and eye position. To compare vision and eye speed contributions, for instance, we computed the vision-observed correlation subtracted by eye speed-observed correlation, in each unit. Along this axis, units with positive values were deemed as vision-driven whereas units with negative values were deemed as eye-speed driven. At the population level along this axis, we examined whether the units with high or low correlation to task performance (AUC > 0.8 versus AUC < 0.8) have the same median value, with Wilcoxon rank-sum test. To assess the extent to which linear visual and eye-movement components explained the neuronal correlate of detection, we first subtracted the model-predicted SDF from the observed SDF derived from spike traces. The discriminability of the neuronal activity for correct versus failed trials was then quantified using the same AUC method applied to the subtracted spikes.

### Neural and Behavioral Latencies

We defined behavioral latency as the time between the target stimulus onset and the saccade onset. To detect saccade onsets, eye position signals were converted to eye speed and acceleration. The times at which the acceleration crossed zero were registered as potential saccade onset times and only recorded if the subsequent eye speed exceeded ten degrees/s. Trials in which the first saccade did not reach the tolerance window for the target stimulus (two-degree radius within target stimulus) were excluded from the analysis.

In each trial, we defined neuronal latency as the time after target stimulus onset when the SDF exceeded a threshold defined as four standard deviations above the mean neural activity from 0 to 500 ms prior to target onset. A neuronal latency was not computed for trials in which the target-evoked response never exceeded the threshold or no spikes occurred. The SDF threshold crossing was required to persist for at least 40 ms. Neuronal latencies were constrained to the interval from 0 to 500 ms after target onset. Thresholds for neuronal latency detection were calculated individually for each trial.

The neural and behavioral latencies were the subject of a population-level analysis. For this analysis, we restricted our analysis to trials in which animals made a saccade to the target stimulus. Units were preselected for inclusion based on the number of trials in which both neuronal and behavioral latencies to the unit’s preferred target direction were reliably detected, with the threshold set at three trials. The relationship between neuronal and behavioral latencies was evaluated using Spearman’s rank correlation. Across the units, we examined distribution of the latency correlation, with respect to their encoding of vision, eye speed and eye position. Along the axis of vision-observed correlation, subtracted by eye-speed-observed correlation, we examined whether the units with significantly high latency correlation (r > 0.5, p < 0.05) and remaining units had the same median value, with Wilcoxon rank-sum test.

## Results

### GLM predicts neural firing during both trained and untrained saccades

The head-fixed animals were exposed to visual stimulation while eye movements were continuously monitored through an infrared camera (**Figure 1A**). Animals were trained to fixate the central fixation point and rewarded for making a saccade to a peripheral target that could appear at one of 8 or 36 locations, with eccentricity five degrees (**Figure 1B**). The peripheral target appeared 700 - 900 ms (M1) or 500 - 700 ms (M2) after fixation was acquired, and only saccades made within 500 ms of the target appearance were rewarded. In some trials, the animals failed to saccade to the target. We analyzed 886 units (828 from M1, 58 from M2) that passed the inclusion criteria (Method) during the task trials as well as in between trials. We examined the neural activity in the period from 0 - 500 ms after appearance of the visual target and found that most neurons were direction tuned (**Figure 1C**), predominately in the visual field contralateral to the recording hemisphere^49^. However, it remains uncertain whether this neuronal firing was driven by the target visual stimuli, or the eye movement or a combination of both.

To investigate how neurons encode the events of visual stimulation and eye movements, we employed an encoding modeling approach. Specifically, we estimated a statistical model that predicts neuronal firing based on events convolved with a linear temporal kernel. We included visual target stimulus, eye speed, eye position, pupil diameter and blinks as explanatory variables, and estimated their linear temporal kernels using ridge regression to predict neuronal firing at each moment during the recording. Notably, the kernels were estimated using data both during and in between task trials.

Neurons in LIP fired both in correct and failed trials, and inter-trial intervals when no visual stimulus was shown. To understand how LIP neurons respond across these different contexts, we conducted event-triggered analysis, exemplified in three neuronal units (**Figure 2A-I**). Specifically, we aligned neuronal firing in three distinct ways: (1) to the onset of the target stimulus for trials in which a saccade was made to the target (**Figure 2A-C**); (2) to the target stimulus for trials in which no saccade was made within the 500 ms response window (**Figure 2D-F**); and (3) to the time of saccade onset for saccades that occurred between trials (**Figure 2G-I**). In all three conditions, neuronal responses to eye movements in the unit’s preferred direction were considered.

**Figure 2.**
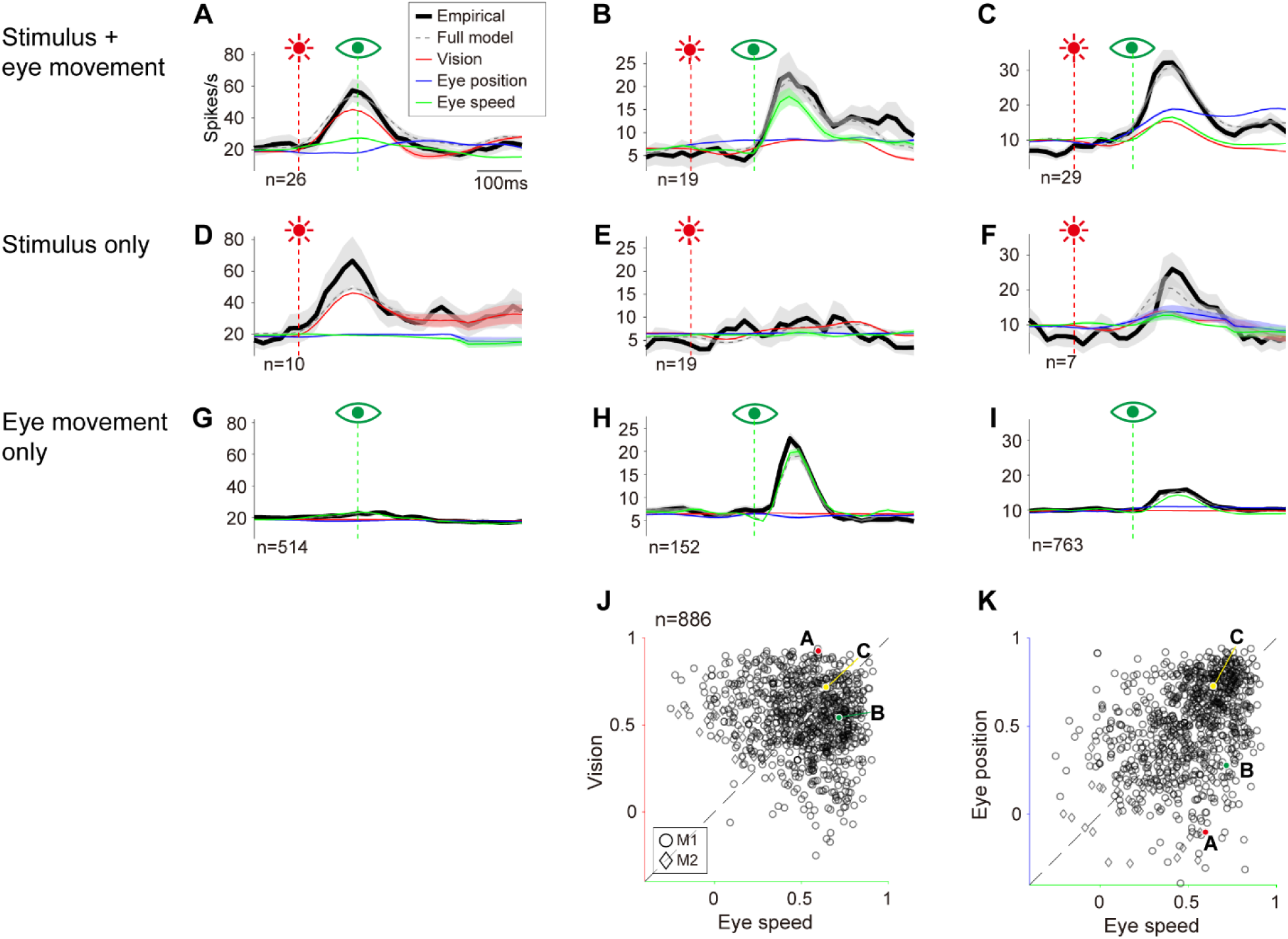
Neural firing is decomposed into stimulus and eye-movement components with GLM. **A-C**. Event-triggered response to the target stimulus presented at the unit’s preferred direction (red vertical line) followed by correct saccade (green vertical line) to the target in three example units. Colored traces in each panel indicate observed firing rate (black), predicted firing rates using the full model (gray), visual kernel (red), eye-speed kernel (green), eye-position kernel (blue). Red, green and blue traces include time-resolved components in addition to the average firing rate over time. **D-F**. Event-triggered response to the target stimulus, without saccade (failed trials). **G-I**, Event-triggered response to the saccade onset in-between trials. **J, K**. Correlation of model components to observed firing 0 - 500 ms after target stimulus onset of correct trials, in vision and eye position space (**J**) and eye position and speed space (**K**). Correlation was computed on trial-averaged traces. In panels **A-I**, n indicates the number of trials or saccades.

All three neurons responded to the target stimulus followed by a saccade (**Figure 2A-C**, *black*), and the GLM accurately captured most features of this response (**Figure 2A-C**, *grey*). At the population level, the GLM provided excellent fits to the observed firing rate during the window from 0 - 500 ms after the target stimulus onset, averaged across correct trials (Pearson correlation = 0.88 ± 0.09, **Figure Supplement 2**). Nevertheless, because this context involves both a visual stimulus and an eye movement occurring closely in time, event-triggered analysis alone cannot determine whether the neurons were responding to the visual stimulus, the eye movements, or a combination of both.

To investigate the drivers of neuronal firing during trials of visual stimulation followed by a correct saccade (**Figure 2A-C**), we utilized the GLM kernels estimated for each unit. Namely, we decomposed observed neuronal activity into components driven by the visual stimulus, eye speed, and eye position, by convolving these explanatory variables with their corresponding GLM kernels. In the first example unit, the response during the correct saccade is primarily driven by the visual stimulus (**Figure 2A**, *red trace*), with minimal contribution from eye movements. In contrast, the second unit’s firing is predominantly influenced by eye position (**Figure 2B**, *green trace*), with little to no contributions from the visual stimulus or eye speed. The third unit displays a mixed response, with contributions from the visual stimulus, eye speed and position components (**Figure 2C**). Notably, this mixed response is unlikely to stem from a result of the predicting GLM, as the ridge regression adopted here should push GLM coefficients towards zero.

To understand how LIP neurons respond to the visual stimulus when no saccade is made, we examined firing during failed trials in which a stimulus appeared, but no saccade was initiated (**Figure 2D-F**). This allows us to isolate the effects of visual stimulation from eye movements. The first and third example units responded in these trials, with their activity largely driven by the visual stimulus component of the GLM (**Figure 2D, F**). This aligns with their responses during correct trials, where the visual stimulus also played a major role (**Figure 2A, C**, *red traces*). Conversely, the second example unit and its model showed negligible response during the failed trials (**Figure 2E**), consistent with its lack of the visual-stimulus driven component. These findings demonstrate that neural responses to the visual stimulus in the absence of saccades vary considerably across units. The GLM effectively captured this unit-specific variability.

We further investigated how LIP neurons fire in between trials, when the animal was not exposed to high-contrast visual stimuli and could move its eyes freely. The first example unit and its model showed no response to saccades in this context (**Figure 2G**), consistent with the absence of eye-movement components in this unit. In contrast, the second and third units exhibited clear responses to the saccades (**Figure 2H, I**), accurately captured by their respective models. Hence, neural response to saccades in the absence of the visual stimulation varies markedly across units, and the GLM effectively captures this unit-specific variability.

Collectively, these analyses demonstrate that neuronal firing in LIP of the marmoset varies markedly depending on the context - whether during visual stimulation with or without saccades, or during spontaneous saccades in between trials, in unit-dependent manner. These contextual variations are succinctly captured by the model estimated by GLM, highlighting the model’s capability to account for the apparent diversity in neural encoding.

To capture the variability in encoding across recorded units, we calculated the correlation across time between the observed firing and the modelled response to correct trials. Correlations were calculated separately for each model component - visual stimulus, eye speed and eye position. This correlation metric indicates the extent to which a unit’s neuronal firing is explainable by the visual stimulus, eye speed or eye position, and is shown in **Figure 2J**, with vision correlation on the y-axis, and the eye-speed correlation on the x-axis. For example, the vision-driven unit (**Figure 2A, D, G**) is positioned in the top-left half, while the eye-speed driven unit (**Figure 2B, E, H**) appears in the bottom-right. The integrator unit (**Figure 2C, F, I**), which contains a combination of visual stimulus, eye speed and position components, is near the identity line. This scatter plot illustrates that the encoding property is broadly distributed rather than tightly clustered into vision-driven and eye-speed driven categories. The correlation metric using eye speed on the x-axis, and eye position on the y-axis is shown in **Figure 2K**. Similar to **Figure 2J**, the integrator unit (**Figure 2C, F, I**) is located near the identity line, along with the majority of other units. These findings indicate that encoding of visual stimulus and eye movements is broadly distributed across neurons in the LIP region.

### Apparent neural correlates of detection are explained by linear kernels

Our analysis has revealed that LIP neurons encode visual stimuli, eye position and speed with considerable variability across neurons. But how does this neural encoding relate to the animal’s behavior? Since LIP neurons are thought to be a part of an inter-regional network responsible for executing visually-guided saccades, we examined two key aspects of saccadic behavior. First, we examined delays between visual stimulation and saccade onset (reaction time). Second, we examined the accuracy of stimulus detection behavior, based on whether a saccade was initiated towards the target stimulus.

How do different types of neural encoding relate to the animal’s detection behavior? To investigate this, we focused on how neural firing might distinguish between trials in which the animal correctly detected the target stimulus and subsequently made a saccade, and trials where it failed to do so. Examining detection-related neural activity requires disentangling motor-related signals from those related to the detection itself. In tasks like the cue-saccade task used in this study, this distinction is challenging due to the overlap between motor-related and detection-related activity. However, the GLM-based modeling approach allows us to effectively "explain away" the motor-related activity by subtracting the convolution of all five explanatory variables with the linear kernels from the observed neuronal firing, leaving behind the component related to the detection. To isolate the detection-related neuronal signal, we limited our analysis to trials in which the target stimulus was presented in the unit’s preferred direction. This ensured that any difference between correct and failed trials were not confounded by variations in the stimulus direction.

We found a sizable fraction of recorded units that appeared to dissociate between the correct and failed trials. One such example is shown in **Figure 3A**, where the unit exhibited vigorous firing about 150 ms after the stimulus onset during the correct trials (**Figure 3A**, *black*), but not during the failed trials (**Figure 3A**, *grey*). In another example shown in **Figure 3B**, the unit fired more in failed trials (**Figure 3B**, *grey*) than in correct trials (**Figure 3B**, *black*) 50 - 400 ms after the visual stimulus onset (AUC = 0.87).

**Figure 3.**
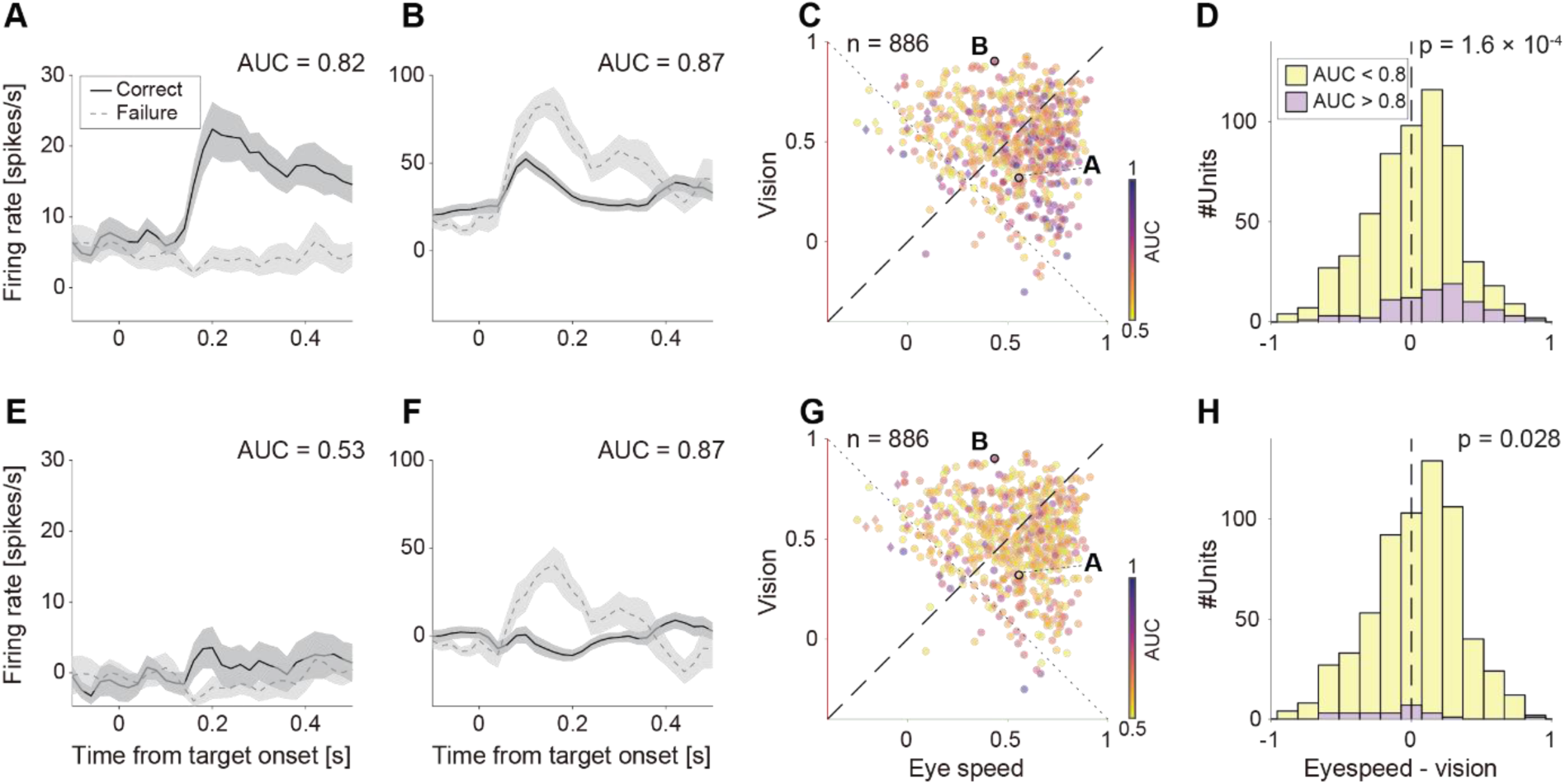
Neural correlates of task performance in observed response (top row) and after regressing away linear components (bottom row). **A, B**. Event-triggered response of two example units, aligned to visual stimulus onset to the preferred direction, followed by correct saccades (black) and without saccades (grey). Top and bottom panels indicate before subtraction of linear components, estimated by the GLM. **C.** Dissociability of correct and failed trials (color-coded) and its relation to encoding of vision (y-axis) and eye-speed (x-axis). The dissociability was measured as each unit’s AUC and represented as a color-coded symbol. Units encircled with black indicate example units displayed in panels **A** and **B**. Units from animals M1 and M2 are indicated with circle and diamond symbols, respectively. **D.** Distribution of units with significant difference between correct and failed trials (AUC > 0.8, purple) and units with insignificant difference (AUC < 0.8, yellow) with respect to their encoding. Here, x-axis indicates relative correlation of eye speed, subtracted by that of visual stimulus, and positive values indicate more preferential encoding of eye speed, and negative values indicate more preferential encoding of visual stimulus. **E-H**, Same as panels **A-D**, after subtraction of the linear components estimated by the GLM.

Can the apparent dissociation between the correct and failed trials be fully explained by the linear components of the GLM? In the first example unit (**Figure 3A**), the dissociation in neural firing between correct and failed trials is indeed almost completely explained away after subtracting the linear components (visual stimulus, eye speed and eye position), estimated by the GLM (**Figure 3E**, AUC = 0.53). This indicates that this example unit is not encoding the detection per se. On the other hand, in the second example unit (**Figure 3B**), the dissociation remained substantial after subtraction of the linear components (**Figure 3F**, AUC = 0.87).

Further analysis of these two examples unit revealed that they primarily encode different aspects of the task. The first example unit (**Figure 3A, E**) primarily encodes eye speed over visual stimulus, because the correlation between the observed firing and the modelled response using the eye-speed kernel is larger than the correlation using the vision kernel (**Figure 3C**). The second example unit (**Figure 3B, F**) primarily encodes visual stimuli, because the correlation between the observed firing and the modelled response using the vision kernel is larger than the correlation using the eye-speed kernel (**Figure 3C**)

At the population level, units with large dissociation between the correct and failed trials (as determined by AUC), were primarily clustered among those driven by eye speed (**Figure 3C**, *purple*). Indeed, the distribution of units with large dissociation between correct and failed trials (**Figure 3D**, *purple*) were more pervasively found in the units that preferentially encode eye-speed, than the units for which correct and failed trials could not be dissociated (**Figure 3D**, *yellow*) (p = 3.3 x 10^-4^, Wilcoxon rank-sum test). However, after subtracting the linear components from the GLM, the ability of eye-speed-driven units to discriminate between correct and failure drastically diminished (**Figure 3G**). As a result, units with high dissociation were distributed more evenly across in vision-driven, eye-speed units together with the integrator units (p = 0.027, Wilcoxon rank-sum test, **Figure 3H)**. A similar trend was observed in the vision and eye-position space (**Figure Supplement 3**).

### Neural latency correlates with behavioral latency in an encoding-type dependent manner

In the previous sections, we focused on trial-averaged spiking rates, which gives no insight into how neural activity and behavior vary across trials. To address this, here we investigate the relationship between single-trial behavioral latency (reaction time) and neural latency. These were quantified as the delay from visual stimulus onset to saccade initiation or the onset of robust neuronal activity. We restricted our analysis to trials in which the animal made a saccade to the stimulus direction within 500 ms, and the firing rate exceeded the four standard deviation threshold. We further restricted the analysis to trials in which the visual stimulus was presented in the neuron’s preferred direction. Although estimating latencies on a single-trial basis yields a relatively small number of trials, this approach enabled us to examine the trial-by-trial variability when the stimulus and saccade directions were aligned.

We identified units whose firing latencies varied systematically with the behavioral latency. An example unit is shown in **Figure 4A1**, where trials were sorted by behavioral latency (*magenta circles*). In this unit, longer behavioral latencies were associated with larger delays in the neuronal firing (*cyan circles*; r = 0.93, p = 1.6 x 10^-3^, **Figure 4A2**). Notably, behavioral latency was consistently about 50 ms shorter than neuronal latency across all trials. That is to say, these neurons’ firing pattern itself cannot causally contribute to saccade initiation, because they are active after, not before onset of the saccade.

**Figure 4.**
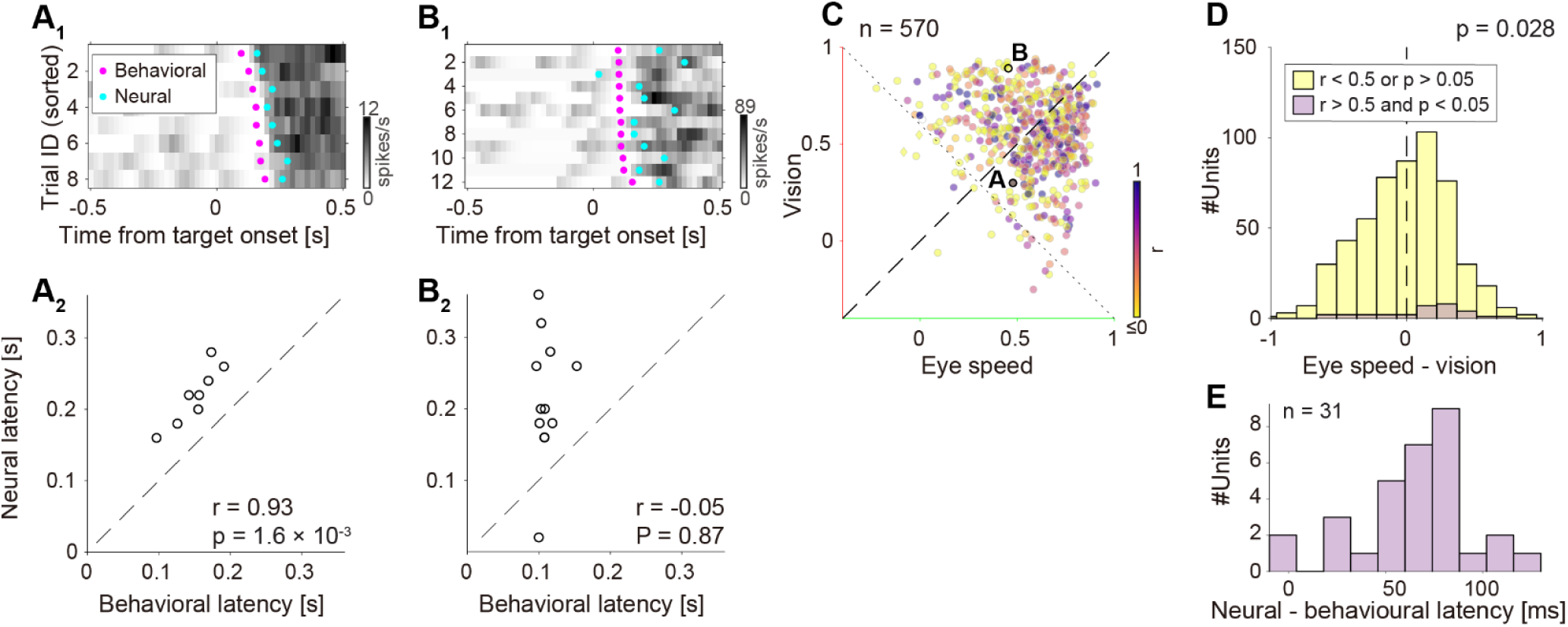
Neural latency correlates with behavioral latency in a cell-type dependent manner. **A, B**. Top: Two example units’ neural firing rate and neural latency (cyan), aligned to the visual stimulus onset to the unit’s preferred direction. Rows are sorted according to behavioral latency (magenta). Bottom: Trial-by-trial correlation (latency correlation) between neural latency (y-axis) against behavioral latency (x-axis). **C**. Variability in latency correlation (color coded) across units, and their dominance in encoding of vision (y-axis) and eye speed (x-axis). Units encircled with black indicate example units displayed in panels **A** and **B**. Units from animals M1 and M2 are indicated with circle and diamond symbols, respectively. **D**. Distribution of units with high latency correlation (> 0.5), with respect to their encoding property. Here, x-axis indicates relative correlation of eye speed, subtracted by that of visual stimulus, and positive values indicate more preferential encoding of eye speed, and negative values indicate more preferential encoding of visual stimulus. **E**. Delay between neuronal latency and behavioral latency in units the latency correlation above 0.5.

We also observed neurons whose firing latencies did not covary with behavioral latency. One such example is shown in **Figure 4B1**, where trials sorted by behavioral latency reveal no consistent relationship with neuronal latency. In this unit, longer behavioral latencies did not correspond to longer neural latencies. Consequently, the behavioral and neuronal latencies were not correlated (**Figure 4B2**, r = -0.05, p = 0.87). This indicates that this unit’s firing pattern does not provide information about the timing of the saccade.

How do these neurons, informative and non-informative about behavioral latencies differ in their encoding of visual and eye-movement signals? To investigate this, we examined how well the visual stimulus and eye-speed component of the model explains the observed firing, measured as the correlation between the modelled response by visual or eye movement components and observed firing. Units that exhibited a strong association between neural and behavioral latencies, such as the example unit in **Figure 4A**, were typically located at the bottom-right half of the plot, indicating a dominant encoding of eye movement (**Figure 4C**, *purple*). Indeed, units whose latency correlation exceeded 0.5 (p < 0.05) were more pervasively found in the bottom-right half, preferentially encoding eye speed rather than visual stimulus (**Figure 4D**, p = 0.028, Wilcoxon rank-sum test). We observed similar results in the vision-eye speed space (**Figure Supplement 4**). Amongst units with high latency correlation, the delay between the neural and behavioral latency had a mean of 72 ms, with neural responses lagging behavioral responses (**Figure 4E**). Together, these findings suggest that neurons that preferentially encode eye movements over visual stimuli are more likely to be informative in predicting saccade latency.

## Discussion

We applied a GLM framework to the relatively understudied domain of rapid, goal-directed behavior to describe the firing patterns of neurons in marmoset LIP. Our approach built on previous applications of GLMs to visual discrimination and memory guided oculomotor tasks in LIP and FEF^31–34^. By incorporating distinct visually-driven and oculomotor components, our model accounts for a substantial portion of neural activity during a rapid, goal-directed saccade task. Critically, the model also accounted for neural activity on trials when saccades were not made, and during the intertrial interval when spontaneous eye movements occurred in the absence of visual stimuli (mean correlation between observed and modeled activity: r = 0.88). We confirmed diverse encoding properties across neurons, with some encoding primarily visual stimuli, some encoding primarily eye-movements, and others encoding both. After demultiplexing these visual and motor components, a negligible proportion of neurons exhibited selective firing correlated with the task performance based on visual detection. Notably, LIP neurons did not exhibit strong pre-saccadic activity but instead showed clear encoding of post-saccadic oculomotor activity, distinct from purely visual activity associated with retinal slip during the saccade. Below, we discuss how these findings align with previous studies in both marmosets and macaques and explore their implications for the functional role of LIP.

*Saccade Initiation.* Pre-saccadic activity has been reported numerous times in macaque^44–46,50^ and in marmoset^15^, and is considered as key evidence that LIP is involved in instantiating saccade-based target selection. However, its precise role remains unsettled. For example, pre-saccadic activity has been shown not to differ between executed and cancelled saccades, raising questions about its causal contribution to saccade initiation^51^. Our investigation revealed post-saccadic neurons but did not identify neurons that consistently fire immediately prior to saccade initiation, particularly under conditions that do not require evidence accumulation for saccade selection. For a neuron to be considered involved in saccade initiation, it must reliably modulate its firing prior to saccade onset. While we did observe neurons whose firing occasionally preceded saccades on individual trials, these responses lacked a consistent, trial-by-trial relationship to the timing of saccade onset. In contrast, neurons whose activity was tightly time-locked to saccade events fired only *after* the onset of the saccade. Previous work in macaques^14^ and marmosets^15^ has reported pre-saccadic activity during gap-saccade tasks, in which a brief delay or “gap” of up to 400 ms is introduced between the disappearance of the fixation target and the appearance of the saccade target. Both studies found relatively little pre-saccadic activity in a saccade task with no gap, as in the task employed in the current study. In macaques, the same cells that showed pre-saccadic activity in the gap-saccade task showed delay period activity in a memory-guided saccade task. In combination with our results, this suggests that pre-saccadic activity is specific to contexts that impose a substantial period to prepare for making a saccade, such as the gap or the memory-guided saccade tasks.

*Encoding of eye position.* Areas of the posterior parietal cortex have also been studied for their role in encoding eye position. Cortical eye position signals are thought to be essential for transforming retinal input, which changes with every eye movement, into a stable visual representation of the world. Neurons in areas of the posterior parietal cortex, including LIP, modulate their firing rates systematically dependent on eye position^3,52,53^. Some neurons appear to shift their receptive fields in anticipation of an upcoming saccade, which may reflect visual remapping^54^. However, despite some neurons showing pre-saccadic modulation in firing rate, eye positions signals are inaccurate until up to 150 ms post-saccade^43^. While we did not explicitly test different eye positions in our current study, a possible explanation for the post-saccadic activity we observed is an updating of eye position signals. *Involvement in Cognitive Process.* In our study, many neurons in marmoset LIP showed modulation depending on whether the animal correctly detected a visual target. However, nearly all of neural modulation occurred after saccades. This finding appears to contrast with a recent report in marmoset LIP^35^, in which the majority of neurons exhibited selectivity for a target stimulus presented alongside a distractor. We propose two potential explanations to reconcile these seemingly conflicting results. First, the task employed in our study required detection of a single supra-threshold target, whereas the task employed in the aforementioned report required discrimination between multiple simultaneously presented stimuli. It is therefore plausible that LIP is not directly involved in target detection per se, as previously suggested^55^, but may play a role in stimulus discrimination when multiple competing stimuli are present. Second, the target selectivity reported previously^35^ may reflect eye movement encoding rather than stimulus discrimination. In their data, neurons that appeared to discriminate targets also showed firing patterns tightly coupled to the direction of the resulting saccade, regardless of distractor presence. This raises the possibility that these neurons are more involved in movement execution than in perceptual selection. To disambiguate these functional roles, demultiplexing analyses, such as the one used in our study, may offer a promising approach for isolating movement-related activity from putative cognitive signals. More broadly, our observation that only a minority of neurons in LIP show evidence of involvement in decision processes aligns with findings from large-scale recordings in mice, where only a small fraction of neurons in any given area are tightly correlated with perceptual decisions^56,57^.

We observed that a large proportion of neurons in marmoset LIP exhibited robust encoding of eye movements, with activity occurring after saccade onset, which are dissociable from visually-evoked activity. This pattern of post-saccadic firing is consistent with previous observations made in macaque^14^ and marmoset LIP^35^. Our results expand on this observation by revealing a tight temporal coupling between saccade onset and neural response, with firing emerging approximately 70 ms after saccade initiation. This temporal delay indicates that these neurons do not contribute to the generation of saccade commands but instead reflect signals occurring after movement onset. Nevertheless, this activity still occurs in the absence of a visual target stimulus, for example when saccades are made during the inter-trial interval, suggesting they play some role in motor control or task monitoring.

We propose three possible interpretations of this post-saccadic activity. First, it may reflect a corollary discharge or efference copy, an internal copy of the saccadic motor command, potentially originating from the SC and conveyed to LIP likely via FEF^58,59^. Second, it could represent motor feedback from oculomotor effectors, conveyed via afferent pathways reporting the outcome of the executed movement. And finally, it could also indicate memorized visual stimulus location, as reported in macaque LIP^48^. Distinguishing between these possibilities will require further targeted investigations, such as the use of the double-step saccade task, which has been instrumental in dissecting corollary discharge circuits in macaques^59^.

As discussed in the sections on saccade initiation and cognitive involvement, the firing patterns of LIP neurons are highly task- and context-dependent. In the present study, using a simple goal-directed saccade task, our modeling approach enabled us to distinguish sensory-from motor-dominated responses. Future work should extend this framework to both simpler and more complex behavioral paradigms, such as the gap saccade task, to clarify how task context shapes the response properties of LIP neurons.

---

**Figure Supplement 2.**
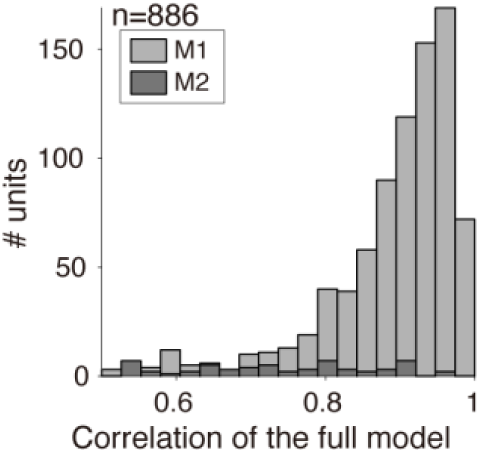
Correlation of the averaged response to the target stimulus across correct trials, between the full model and the observed firing rate traces.

**Figure Supplement 3.**
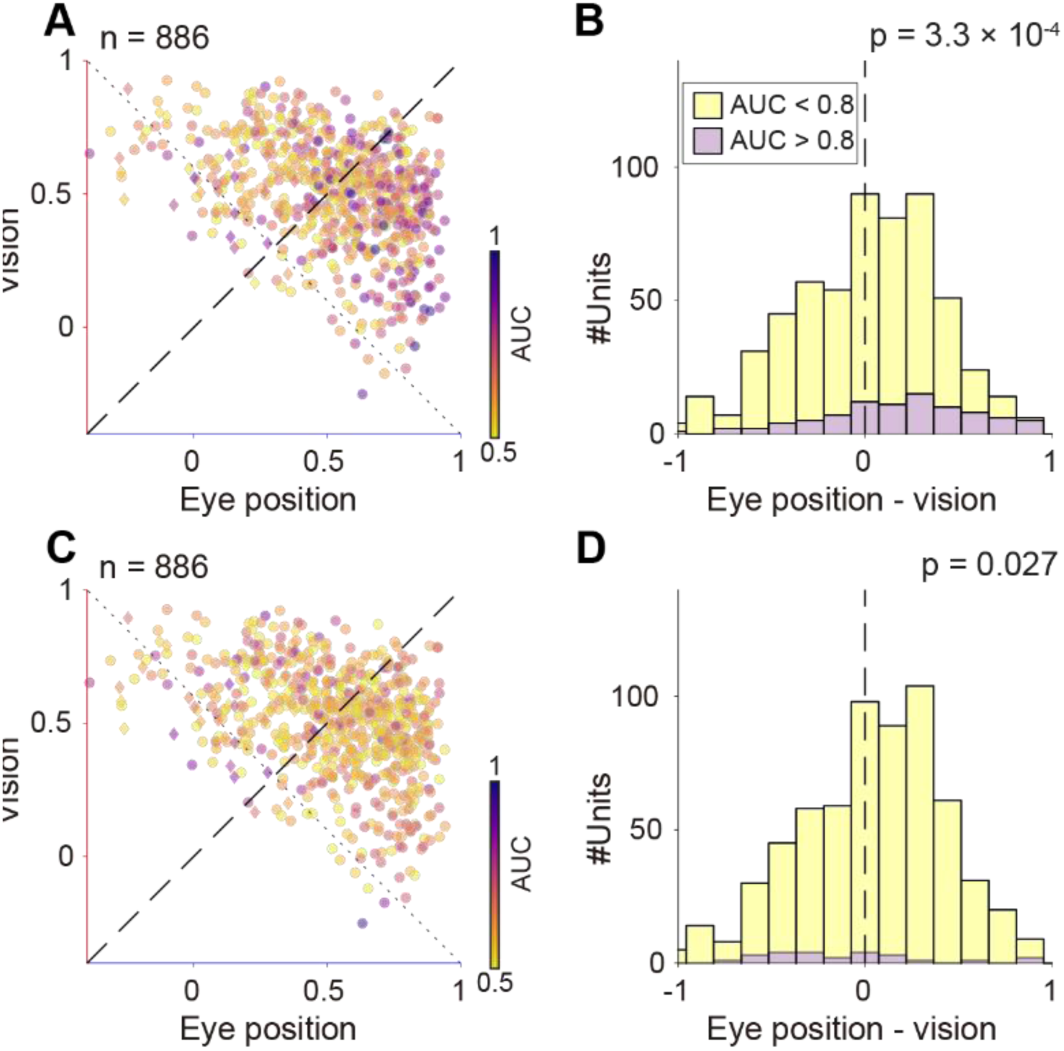
**A**. Dissociability of correct and failed trials (color-coded) and its relation to encoding of vision and eye-position. Units from animals M1 and M2 are indicated with circle and diamond symbols, respectively. **B**. Distribution of units with significant difference between correct and failed trials (AUC > 0.8, purple) and units with insignificant difference (AUC < 0.8, yellow) with respect to their encoding. Here, x-axis indicates relative correlation of eye position, subtracted by that of visual stimulus, and positive values indicate more preferential encoding of eye speed, and negative values indicate more preferential encoding of visual stimulus. **C, D**, Same as panels **A, B**, after subtraction of the linear components estimated by the GLM.

**Figure Supplement 4.**
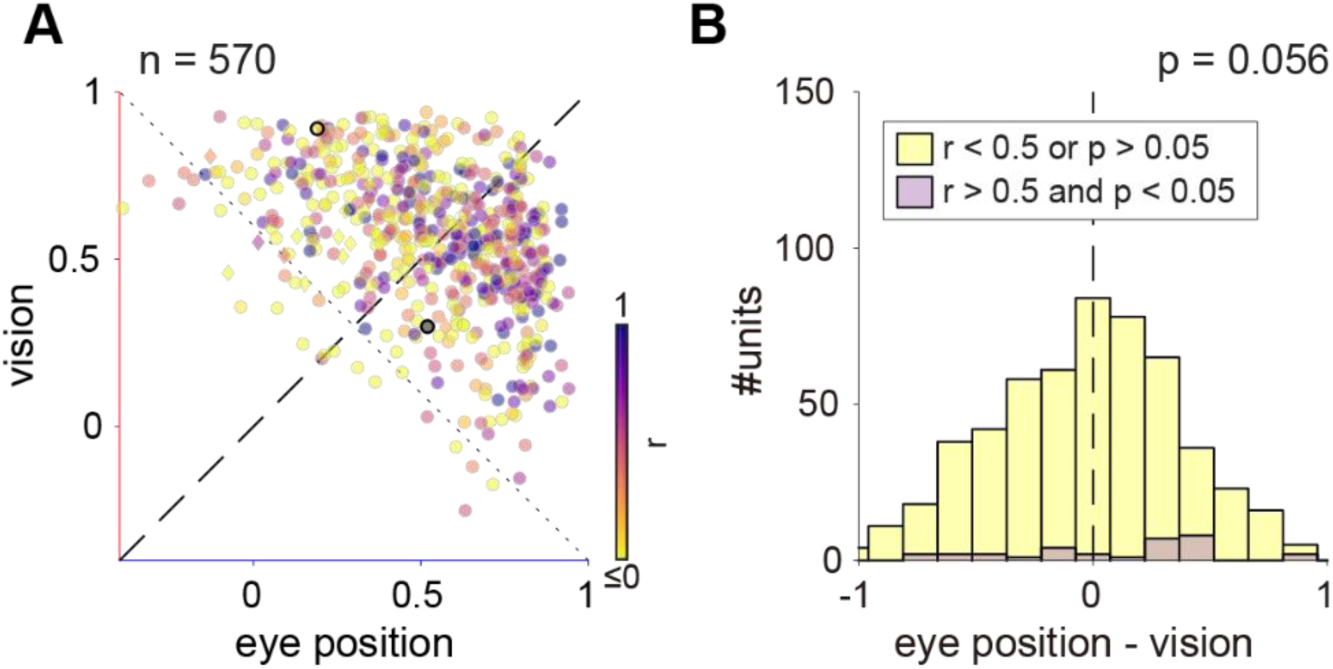
**A**. Neural latency correlates with behavioral latency in vision-eye position space. Units from animals M1 and M2 are indicated with circle and diamond symbols, respectively. **B**. Distribution of units with high latency correlation (> 0.5), with respect to their encoding property. Here, x-axis indicates relative correlation of eye position, subtracted by that of visual stimulus.

